# Neural differentiation of incorrectly predicted memories

**DOI:** 10.1101/083022

**Authors:** Ghootae Kim, Kenneth A. Norman, Nicholas B. Turk-Browne

## Abstract

When an item is predicted in a particular context but the prediction is violated, memory for that item is weakened (Kim et al., 2014). Here we explore what happens when such previously mispredicted items are later re-encountered. According to prior neural network simulations, this sequence of events - misprediction and subsequent restudy - should lead to differentiation of the item's neural representation from the previous context (on which the misprediction was based). Specifically, misprediction weakens connections in the representation to features shared with the previous context, and restudy allows new features to be incorporated into the representation that are not shared with the previous context. This cycle of misprediction and restudy should have the net effect of moving the item‘s neural representation away from the neural representation of the previous context. We tested this hypothesis using fMRI, by tracking changes in item-specific BOLD activity patterns in the hippocampus, a key structure for representing memories and generating predictions. In left CA2/3/DG, we found greater neural differentiation for items that were repeatedly mispredicted and restudied compared to items from a control condition that was identical except without misprediction. We also measured prediction strength in a trial-by-trial fashion and found that greater misprediction for an item led to more differentiation, further supporting our hypothesis. Thus, the consequences of prediction error go beyond memory weakening: If the mispredicted item is restudied, the brain adaptively differentiates its memory representation to improve the accuracy of subsequent predictions and to shield it from further weakening.

**Significance:** Competition between overlapping memories leads to weakening of non-target memories over time, making it easier to access target memories. However, a non-target memory in one context might become a target memory in another context. How do such memories get re-strengthened without increasing competition again? Computational models suggest that the brain handles this by reducing neural connections to the previous context and adding connections to new features that were not part of the previous context. The result is neural differentiation away from the previous context. Here provide support for this theory, using fMRI to track neural representations of individual memories in the hippocampus and how they change based on learning.

## Introduction

When faced with a familiar situation, we can often predict who or what will appear. What happens to the memories supporting these predictions when they turn out to be wrong? We previously found that incorrect prediction of an item in a familiar context leads to worse subsequent memory for that item (Kim et al., 2014). Through this process, the brain might prune memories that correspond to changed or unstable aspects of the environment. However, an item that is irrelevant in one situation might later become relevant in another situation. In this case, the previously weakened memory needs to re-gain its mnemonic strength. How does the brain accomplish this goal? Based on our previous network modeling work (Norman et al., 2006, 2007), and empirical findings (Hulbert & Norman, 2015), we propose that the brain resolves this by adaptively *differentiating* the memory from its previous context.

The model’s predictions are illustrated in Figure 1. Consider two items, A and B, that have been paired with each other previously, such that the appearance of A leads to a prediction of B. But, on this particular trial, B does not appear. Memory A is strongly activated (because it was just shown) and memory B is moderately activated (because it is being predicted from memory). A key postulate in the model is that moderate activation weakens connections from other, strongly activated features (Norman et al., 2006, 2007). Thus, connections from the strongly activated features of A to the moderately activated features of B (those that are not shared with A) are weakened, effectively “shearing” these unique features of B from features shared with A. This weakening of connections within the B representation is a possible explanation for our previous findings of impaired recognition of mispredicted items (Kim et al., 2014). Crucially, if B is restudied later, the unique features of B will be activated, but not features previously shared with A (because connections to these features were weakened). Instead, activation will spread to other new features (not previously shared with A), and connections to these features will be strengthened. This process of swapping out shared for unshared features decreases overlap between the A and B memories.

**Figure 1.**
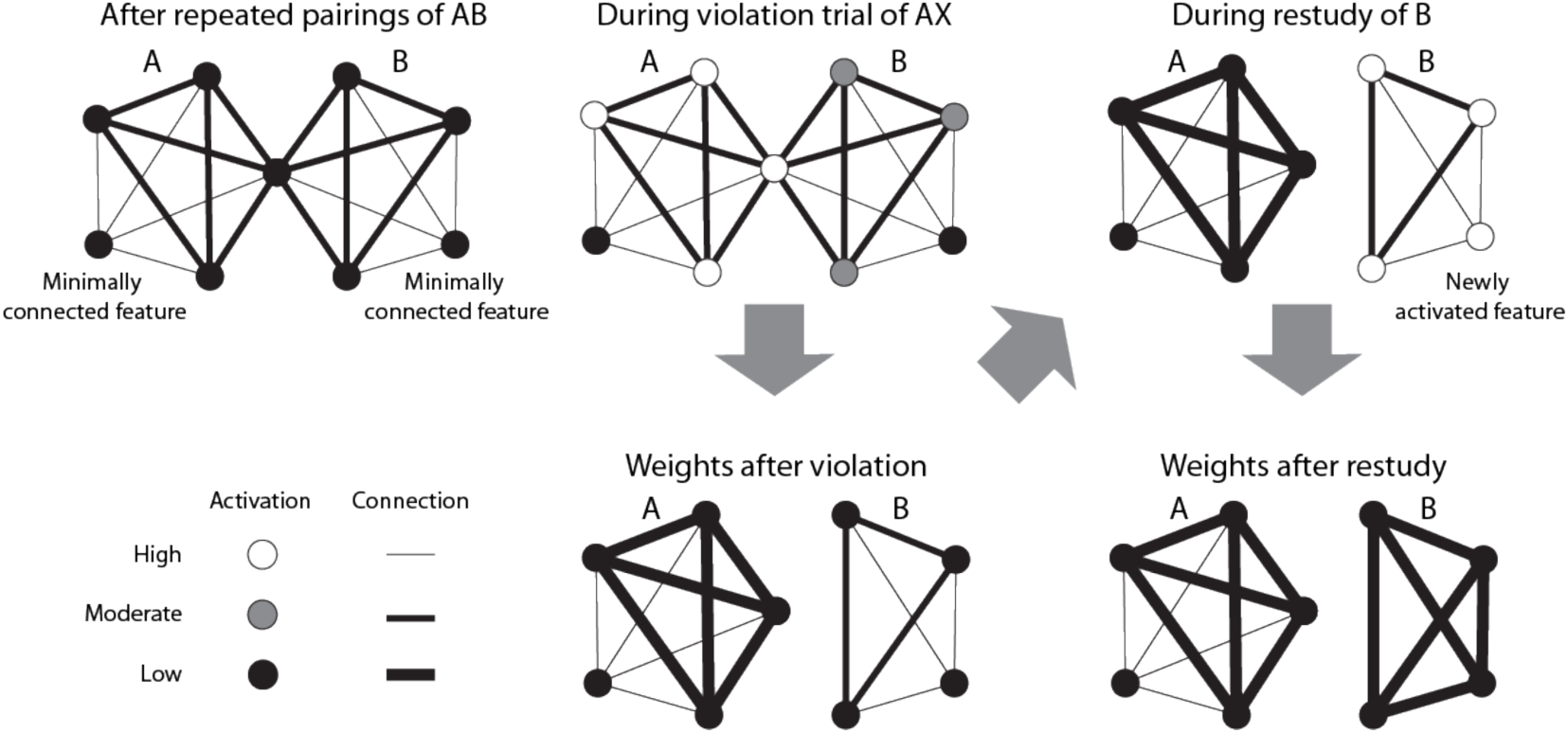
How interleaved misprediction and restudy lead to differentiation. A and B have been paired previously (AB), but in this instance of A, B does not appear (AX). This unconfirmed prediction leads to moderate activation of the features of B. According to our theory (Norman et al., 2006, 2007; Hulbert & Norman, 2015), this leads to weakening of connections into these moderately active features from other, strongly activated features (including features shared with A). If the B item is restudied later after a novel item, activation spreads to new features that were not formerly activated by A, resulting in strengthening of these connections to new features. This cycle — whereby misprediction of B causes shearing of B features from A features, and restudy leads to acquisition of new features — has the overall effect of differentiating B’s neural representation from A’s neural representation.

Other studies have demonstrated neural differentiation from learning of inter-related materials (Favila et al., 2016; Schapiro et al., 2012; Schlichting et al., 2015). The key contribution of the present work is that we provide a mechanism for differentiation (described above). Our proposed mechanism leads to two specific, testable claims that go beyond basic differentiation: First, the mispredicted item should specifically differentiate from its prior context (as opposed to becoming generally more distinct from other items). Second, across items, the degree of misprediction should relate to the degree of differentiation (insofar as misprediction leads to shearing off of shared features, opening the door for new features to be acquired).

We tested these two hypotheses in an fMRI study. Observers were exposed to a continuous sequence of scenes and faces while performing a cover task. Unbeknownst to them, this sequence was generated from pairs (e.g., scene A-scene B), creating an expectation that B will follow A. For some pairs, these expectations were violated (A was followed by X instead of B). All B items were subsequently restudied. We hypothesized that misprediction of B followed by its restudy would lead to differentiation of B from A, compared to a control condition consisting of pairs where expectations were not violated. To test this, we used fMRI and pattern similarity analysis to track changes in neural overlap between A and B and to track how strongly B was predicted on violation trials. Our results show, a direct relationship between competitive dynamics (i.e., misprediction) during learning and representational change, thereby supporting for our mechanistic model of memory updating.

## Materials and Methods

### Participants

Thirty-two adults (19 women, 27 right-handed, mean age 20.1 years) participated for monetary compensation. All had normal or corrected-to-normal vision and provided informed consent. The Princeton University IRB approved the study protocol.

### Stimuli

Participants were shown color photographs of indoor and outdoor scenes (including from http://cvcl.mit.edu/MM/sceneCategories.html), male and female faces (including from www.macbrain.org/resources.htm), and natural and manmade objects. Stimuli were projected on a screen behind the scanner, viewed with a mirror on the head coil (subtending 8.8 × 8.8°). Participants fixated a black central dot that changed to white when a response was recorded.

### Procedure

The experimental procedure unfolded over two days (Fig. 2A). All phases of the experiment were scanned with fMRI. The first session consisted of six runs of an incidental encoding task. Participants viewed face and scene stimuli one at a time and made male/female judgments for faces and indoor/outdoor judgments for scenes. Unbeknownst to participants, the stimulus sequence contained image pairs (e.g., scene A-scene B). There were eight pairs for each of the two conditions (violation and non-violation) within each run. The first and second members of each pair were presented once separately (randomly intermixed with items from other pairs), before they were ever shown together in a pair. This allowed us to measure the neural representation of each item on its own before learning (“pre-learning snapshot”), uncontaminated by the pairmate (Schapiro et al., 2012).

There was no explicit distinction between the pre-learning snapshots and the presentation of pairs. After the items in a given pair were shown separately (to acquire pre-learning snapshots), the items were shown as a pair three times, interleaved with repetitions of other pairs. For pairs assigned to the violation condition, the three repetitions were followed by a violation event (e.g., scene A-face X); this event was omitted for non-violation pairs. Crucially, in both conditions, the B item was subsequently presented (“restudied”) on its own, following a novel item in the sequence rather than its previous context A. This cycle of violation and restudy was repeated once more for the violation pairs — our modeling work suggested that two cycles would produce more differentiation than one — leading to the following overall event sequence per pair: AB-AB-AB-AX-B-AY-B. There was also a second restudy for the non-violation pairs, leading to a sequence that was matched for the number of exposures to B, but without violation events: AB-AB-AB-B-B. Importantly, these manipulations were incidental to the primary task of making categorical judgments, and the interleaving of multiple pairs obscured the pair structure. In a separate behavioral pilot, we encouraged participants to report any regularity, and none noticed that the sequence was constructed from repeated pairs. Each trial began with a blink of the fixation cross to signal an upcoming stimulus, followed by the stimulus presentation for 1 s and a blank interval of 2 s. There were a total of 192 trials (32 pre-learning, 160 pair sequences) within each run, which lasted 10 min and 6 s.

The second session occurred the day after the first session (at least 12 hours) and consisted of three tasks: post-learning snapshot (2 runs), surprise memory test (2 runs), and functional localizer (3 runs). The post-learning snapshots were collected in the same manner as the pre-learning snapshots: all scenes from the first session were shown again, one at a time, in a random order, and with indoor/outdoor judgments. In the recognition memory test, we presented each participant with all B scenes from both conditions (48 violation and 48 non-violation), randomly intermixed with 48 novel scenes. A two-step memory response was made for each scene: “old” or “new”, then confidence level (“sure” or “unsure”). Images remained on the screen until the responses were made, and the next trial began on the first subsequent TR.

The four post-learning and memory runs were interleaved and their order was counterbalanced across participants (odd participants: memory run 1, post-learning run 1, post-learning run 2, memory run 2; even participants: post-learning run 1, memory run 1, memory run 2, post-learning run 2). Half of the studied pairs were assigned to memory run 1 and post-learning run 1, and the other half to memory run 2 and post-learning run 2. In other words, for half of the B items from the first session, memory was tested before the post-learning snapshot was taken, and for the other half of B items, the post-learning snapshot was taken before memory was tested. We designed the procedure this way because we were concerned that testing memory for an item could contaminate the subsequent post-learning snapshot for that item, and vice versa. We wanted some items to get an uncontaminated post-learning snapshot (before memory was tested). Although behavioral recognition memory performance was not the focus of this paper, we also wanted some items to get an uncontaminated memory judgment (before the snapshot was taken). For the analyses focusing on neural differentiation (and related follow-up analyses), we used only the pairs for which the post-learning snapshot preceded the recognition memory test, thereby ensuring that the post-learning snapshot would not be affected by any learning that occurred as a result of recognition memory test.

After the post-learning and memory runs, participants completed three runs of a functional localizer. Each run contained 15 blocks, with five blocks from each of three categories: faces, scenes, and objects. Participants judged faces as male or female, scenes as indoor or outdoor, and objects as manmade or natural. Each stimulus was presented for 500 ms, followed by a blank interval of 1000 ms. There were 10 trials per block and each 15-s block was followed by 15 s of fixation, treated as a rest category. The duration of each run was 465 s. For the analyses below, we used only the scene blocks. Specifically, we calculated a “template” activity pattern for the scene category from the scene blocks; this template was later used to evaluate whether our results were item-specific.

**Figure 2.**
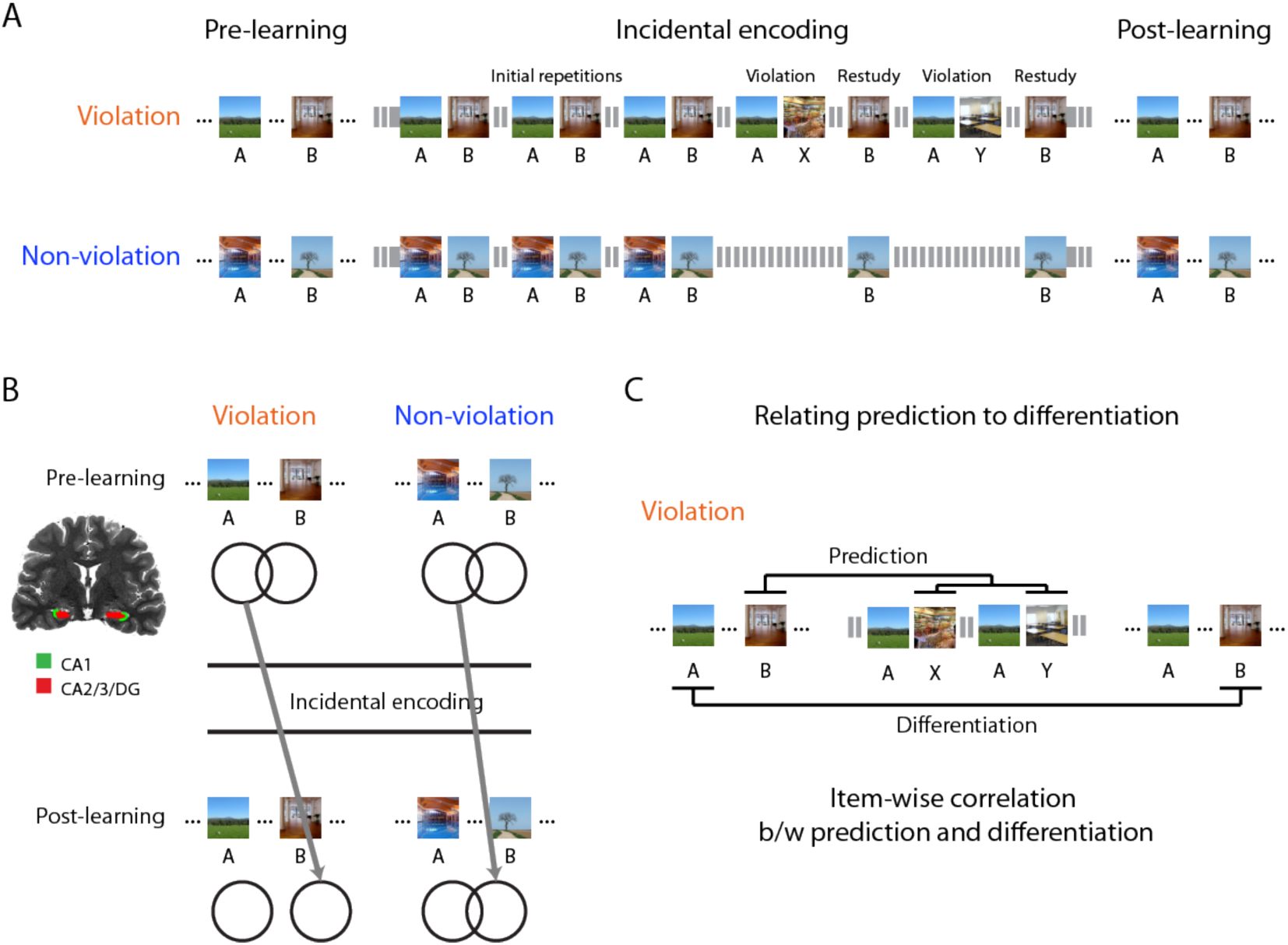
Experimental design and main analyses (A) During incidental encoding, participants performed a categorization task for scenes (indoor/outdoor) and faces (male/female). Before the main incidental encoding phase, all scene images were presented in a random order so that we could take pre-learning snapshots of the neural representations of these images. During the incidental encoding phase, the trial sequence for the violation condition was constructed from three initial repetitions of AB pairs (AB-AB-AB), and two cycles of violation and restudy trials (AX-B-AY-B), whereas the violation trials were omitted for the control (non-violation) condition (AB-AB-AB-B-B). Each individual trial in these sequences was interleaved with trials of other pairs, at different points in their sequences. After the main incidental encoding phase, the same scene images were presented again in a random order, allowing us to take post-learning snapshots of their neural representations. (B) In left CA2/3/DG, we measured neural differentiation by computing the similarity of the pre-learning snapshot of A and the post-learning snapshot of the corresponding B item, in the violation condition vs. the control condition. (C) We measured prediction strength in the violation condition by computing the similarity of the pre-learning snapshot of B to neural patterns present at the moment of the violation (when X/Y items were presented). We then computed the correlation between prediction strength and neural differentiation in a trial-by-trial fashion.

### Data acquisition

The experiment was programmed in Matlab using the Psychophysics Toolbox (http://psychtoolbox.org). MRI data were acquired with a 3T MRI scanner (Siemens Prisma) with a 64-channel head coil. Functional scans came from a multi-band EPI sequence (TR = 1.5 s, TE = 39 ms, flip angle = 50°, voxel size = 1.5 mm iso), with 52 oblique axial slices (transverse to the long axis of the hippocampus) in an interleaved order. A whole-brain T1 MPRAGE image and a co-planar T1 FLASH image were acquired for registration to other participants. Two T2 TSE images were acquired for probabilistic segmentation of hippocampal subfields (54 slices perpendicular to the long axis of the hippocampus; 0.44 x 0.44 mm in-plane, 1.8 mm thick). A field map was acquired to correct for B0 inhomogeneity.

### Regions of interest

This study involves rapidly learning new, arbitrary associations between stimuli. As the hippocampus supports the encoding and retrieval of such memories (Davachi, 2006; Norman & O’Reilly, 2003; Schapiro et al., in press), we expected that representational changes would occur in the hippocampus. Based on a previous study of neural differentiation (Hulbert & Norman, 2015), we focused specifically on the left hippocampus. Within the hippocampus, we were primarily interested in the CA3 and dentate gyrus (DG) subfields, as they are core storage sites for episodic memories and generate predictions via pattern completion (e.g., Hindy et al., 2016).

### Hippocampal segmentation

Subfields of the hippocampus (CA1 and CA2/3/DG) were defined probabilistically in MNI space based on a database of manual hippocampal segmentations from a separate set of 24 participants. Manual segmentations were created on T2-weighted TSE images using anatomical landmarks (Duvernoy, 2005; Carr et al., 2010; Aly & Turk-Browne, 2016), and then registered to an MNI template. Voxels in the MNI template were assigned subfield labels if the probability was greater than 0.5. Nonlinear registration (FNIRT; Andersson et al. 2007) was used to register each participant’s high-resolution MPRAGE to this probabilistic label atlas. Subfields were extracted separately from each hemisphere, and merged for bilateral analyses.

### Preprocessing

fMRI data were preprocessed with FSL (http://fsl.fmrib.ox.ac.uk). Functional scans were corrected for slice-acquisition time and head motion, high-pass filtered (128 s period cut-off), and aligned to the middle volume within each run. As a second alignment step, all preprocessed images in the first session were aligned to the first volume of the first functional run. Functional scans from the second session were aligned to the same volume, based on first aligning the MPRAGE scans across sessions.

### Measuring differentiation

For our initial analysis of overall differentiation, we measured how much B’s neural representation after learning moved away from the original representation of A, and whether this differed for violation and non-violation pairs. Specifically, we measured the Pearson correlation between the pre-learning snapshot of A and the post-learning snapshot of B (Fig. 2B). These snapshots were defined by the spatial pattern of activity elicited by each item in a particular ROI (e.g., left CA2/3/DG), at the peak of the hemodynamic response (4.5 s after image onset). This approach differs slightly from previous studies of representational change that used similar neural snapshot methods (Favila et al., 2016; Schapiro et al., 2012). For example, Favila et al. (2016) measured neural overlap between competing items *within* a post-learning phase, relative to non-competing control items. Such an approach was possible because there was no difference between items in that phase other than their learning history. However, in our study, A in the violation condition was presented two times more than A in the non-violation condition (because of AX and AY violation trials), so any comparison between violation and non-violation conditions that includes the post-learning snapshot of A is confounded by item frequency. Consequently, we used the pre-learning snapshot of A (prior to any difference between conditions) as the baseline for representational change.

Our hypothesis posits that differentiation effects should be item-specific: Weakening of connections between the shared features of A/B and the unique features of B (as a result of misprediction) allows for the subsequent addition of new features to B when it is restudied, and this leads to an overall decrease in neural overlap between A and B. In other words, it is important for our hypothesis that B become more distinct from A specifically, and not just generically more distinct from other items. The basic measure of differentiation above is consistent with both possibilities, and thus we performed an additional randomization analysis to verify item specificity: We scrambled the pair assignments of A and B 1,000 times within each condition and re-calculated neural differentiation. That is, if A_i_ and B_i_ indicate that these items were from the same pair (i), the original analysis involves calculating differentiation between A_1_ and B_1_, A_2_ and B_2_, A_3_ and B_3_, etc., whereas a given permutation in the randomization test might compare A_1_ and B_7_, A_2_ and B_4_, A_3_ and B_2_, etc. If differentiation occurs in a generic sense, then the A items are interchangeable and the original effect will not differ from the permuted distribution. If, as predicted by our model, differentiation is item-specific, the original magnitude of differentiation should be larger than the permuted distribution. For statistical analysis, within each participant, we measured the difference in pattern similarity of pre-learning A to post-learning B between the violation and non-violation conditions for both the correct pairing and the permuted pairings, and computed the *z*-score of the true difference relative to the 1,000 permuted differences. We then examined the reliability of these *z*-scores across participants with a one-sample t-test against zero.

### Relating prediction to differentiation

Beyond showing item-specific neural differentiation, a key contribution of our study is to provide an explanation for how it arises — as a result of misprediction. The main effect of differentiation for the violation vs. non-violation conditions is consistent with this, as they only differed in terms of the presence of violation trials that allowed for misprediction. Nevertheless, we sought stronger and more continuous evidence by attempting to relate, on a trial-by-trial basis, the amount of prediction on violation trials to the amount of subsequent differentiation. This analysis was performed only for the violation condition because there were no violation trials in the non-violation condition. To measure prediction on violation trials, we calculated the amount of evidence for B during the presentation of unexpected items X and Y (i.e., the items that appeared after A, when B should have been presented). Specifically, we measured pattern similarity between the pre-learning snapshot of B (not used to calculate differentiation) and the pattern of activity evoked by both X and Y events, and then averaged these two similarities to provide a single index of prediction for each AB pair. Across pairs, we then calculated the correlation of this prediction score with the pair-specific neural differentiation effect within participant, and then examined its reliability at the group level using a one-sample t-test.

Again, our hypothesis about a relationship between prediction strength and differentiation is item-specific — the activation of B in particular is what induces competition with, and differentiation from, A. However, the analysis above does not guarantee that the correlation is item-specific: Pattern similarity between the pre-learning snapshot of B and X/Y could reflect prediction of item-specific features of B or more generic categorical features of B shared across items (i.e., predicting that a scene will appear, without specifying the scene). We conducted two further analyses to address this. First, we performed the same type of randomization test used above for the main effect of differentiation. Specifically, if prediction reflects generic scene activation, then the pre-learning snapshots of different B items should be interchangeable when calculating pattern similarity with x and Y. Thus, we scrambled the original pairings of pre-snapshot B and X/Y 1,000 times, re-calculated their pattern similarity, and then re-computed the prediction-differentiation relationship. For example, if the original prediction scores were derived by correlating pre-B_1_ with X_1_/Y_1_ (where X_1_ and Y_1_ followed A_1_), and the original differentiation scores were derived by correlating pre-A_1_ with post-B_1_, a permutation might involve re-computing the prediction scores (comparing pre-B_7_ to X_1_/Y_1_, etc.) but keeping the differentiation scores the same; the re-computed prediction scores were then correlated with the differentiation scores, yielding a new (null) value for the prediction-differentiation relationship. As before, a *z*-score for the original prediction-differentiation relationship was calculated relative to the permuted distribution for this relationship within participant, and these *z*-scores were assessed for reliability with a one-sample t-test across participants. According to our hypothesis (but not a generic scene prediction account), permuting the items in this way should abolish the relationship between prediction and differentiation.

Second, we used regression to remove generic category-level information from the activity patterns prior to calculating pattern similarity. Specifically, we defined a template activity pattern for the scene category by averaging over the many scene images in the localizer; we then regressed this template separately from each of the patterns used for this analysis (i.e., we scaled the scene template to maximize fit to the observed pattern and then took the residuals). By definition, the residual patterns after this regression are orthogonal to the scene template, thereby reducing the possibility that generic scene prediction drove the prediction score. As the final step, we repeated the original prediction-differentiation analysis with these residuals. According to our hypothesis, the relationship should be preserved.

## Results

### Behavioral performance

Judgments in the categorization cover task (indoor/outdoor for scenes, male/female for faces) were accurate in general (mean = 90.51%, SD = 10.91; vs. chance: *t*(31) = 21.00, *p* <0.001), and did not differ across the pre-learning snapshots, pair sequences, or post-learning snapshots (*F* < 1). For the recognition memory test, we restricted analysis to the B items that were tested before the post-learning snapshot was taken. Performance was good in general, with B items more likely to be endorsed as “old” than new items (mean *A'* = 0.84, *t*(31) = 26.17, *p* < 0.001). We did not have a specific expectation for a difference in memory between violation and non-violation conditions because neural differentiation can have opposite effects on the underlying processes that support recognition memory (see Discussion). Indeed, there was no difference between conditions (*t*(31) = −0.24, *p* = 0.81). Neural analyses were restricted to the other B items, whose post-learning snapshots were collected prior to the (potentially contaminating) memory test.

### Neural differentiation

We examined how much the neural representation of the B items moved away from their initial A context items by measuring pattern similarity between the post-learning snapshot of B and the pre-learning snapshot of A. We focused initially on left CA2/3/DG ROI, given prior findings of differentiation (Hulbert & Norman, 2015). We hypothesized that misprediction of B items in the violation condition would reduce subsequent neural overlap with A after restudy. Indeed, pattern similarity was lower for pairs from the violation vs. non-violation conditions (*t*(31) = −2.82, *p* = 0.008; Fig. 3A). We conducted additional exploratory analyses outside of the main ROI, in other potentially relevant hippocampal subfields: right CA2/3/DG, and left and right CA1. None of these regions showed a reliable difference between conditions, and right CA1 actually showed a trend in the opposite direction from left CA2/3/DG (left CA1: *t*(31) = 0.31, *p* = 0.76; right CA1: *t*(31) = 1.87, *p* = 0.07; right CA2/3/DG: *t*(31) = 0.69, *p* = 0.50).

Our hypothesis explains this neural differentiation in terms of swapping out shared features with A for new features of B. Thus, the representational change should be specific to B’s relationship with A, and not reflective of increased distinctiveness from other items in general. We evaluated this possibility by permuting the relationship between the pre-learning snapshots of A and the post-learning snapshots of B (Fig. 3B). If neural differentiation is item-specific, then this scrambling should eliminate the difference in pattern similarity between violation and non-violation conditions. Indeed, consistent with our model, the differentiation effect in left CA2/3/DG was stronger for the correct AB pairings than the null distribution of permuted pairings (*t*(31) = −2.81, *p* = 0.009; Fig. 3C).

**Figure 3.**
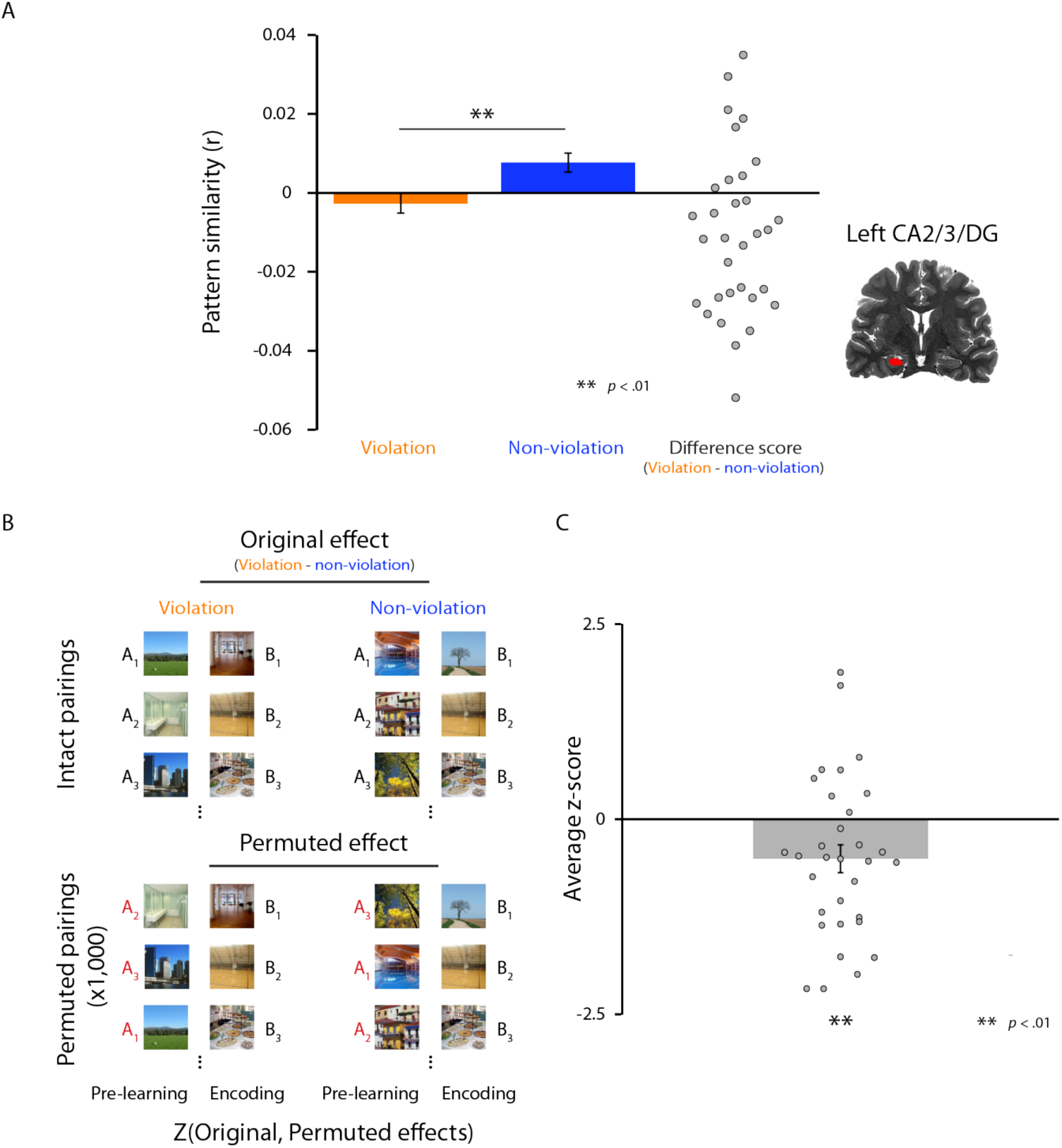
Neural differentiation effect and item-specificity test in left CA2/3/DG. (A) Pattern similarity between the pre-learning snapshot of A and the post-learning snapshot of B was significantly lower for the violation vs. non-violation conditions. (B) Schematic of procedure for randomization analysis. For each participant, we shuffled the AB pairings 1,000 times within each condition. In each iteration, we calculated the same pre-learning A to post-learning B pattern similarity for each condition and stored the difference between conditions. This produced a null distribution of differences, and we calculated the *z* score of the original effect with respect to this distribution. The reliability of the *z*-scores was assessed across participants. (C) Pattern similarity was lower for the original vs. permuted AB pairings, consistent with the neural differentiation effect being item-specific.

### Relationship between prediction and differentiation

The analysis above relies on the categorical manipulation of condition (violation vs. non-violation) to establish that misprediction is responsible for differentiation upon restudy. To provide further support for this hypothesized mechanism, we conducted continuous analyses within the violation condition. According to our model, greater prediction of B during the AX/AY violation trials induces more weakening of its shared features with A, which in turn allows for better subsequent acquisition of new features during restudy of B in a novel context. Accordingly, in left CA2/3/DG, there should be a negative relationship between the amount of prediction of B on violation trials (pattern similarity of X/Y with the pre-learning B snapshot) and the strength of neural overlap for the corresponding pair (pattern similarity between pre-learning A and post-learning B snapshots). Indeed, as shown in Fig. 4A, there was a reliable negative correlation between these measures (*t*(31) = −2.60, *p* = 0.01).

As with the differentiation effect, our model also posits that prediction of B *per se* is critical, as more generic prediction (e.g., of a scene) would not specifically weaken the features of B. We evaluated this possibility using two different analyses: First, we performed a randomization test by permuting the pre-learning B snapshots when calculating prediction during violation trials (under the null hypothesis that the B items are interchangeable) and then recomputed the trial-by-trial relationship with differentiation (Fig. 4B). Consistent with item-specific prediction being the critical ingredient, the relationship between prediction strength and differentiation was stronger for the original pairings of B items and violation trials, relative to the null distribution from permuted pairings (*t*(31) = −2.55, *p* = 0.02; Fig. 4C). Second, we regressed generic category-level information out of the pre-learning B snapshots and violation trials prior to calculating prediction scores (thus attenuating the contribution of non-item-specific features of B to pattern similarity) and then we re-ran the analysis. As expected, the negative relationship between prediction and differentiation persisted (*t*(31) = −2.45, *p* = 0.02).

**Figure 4.**
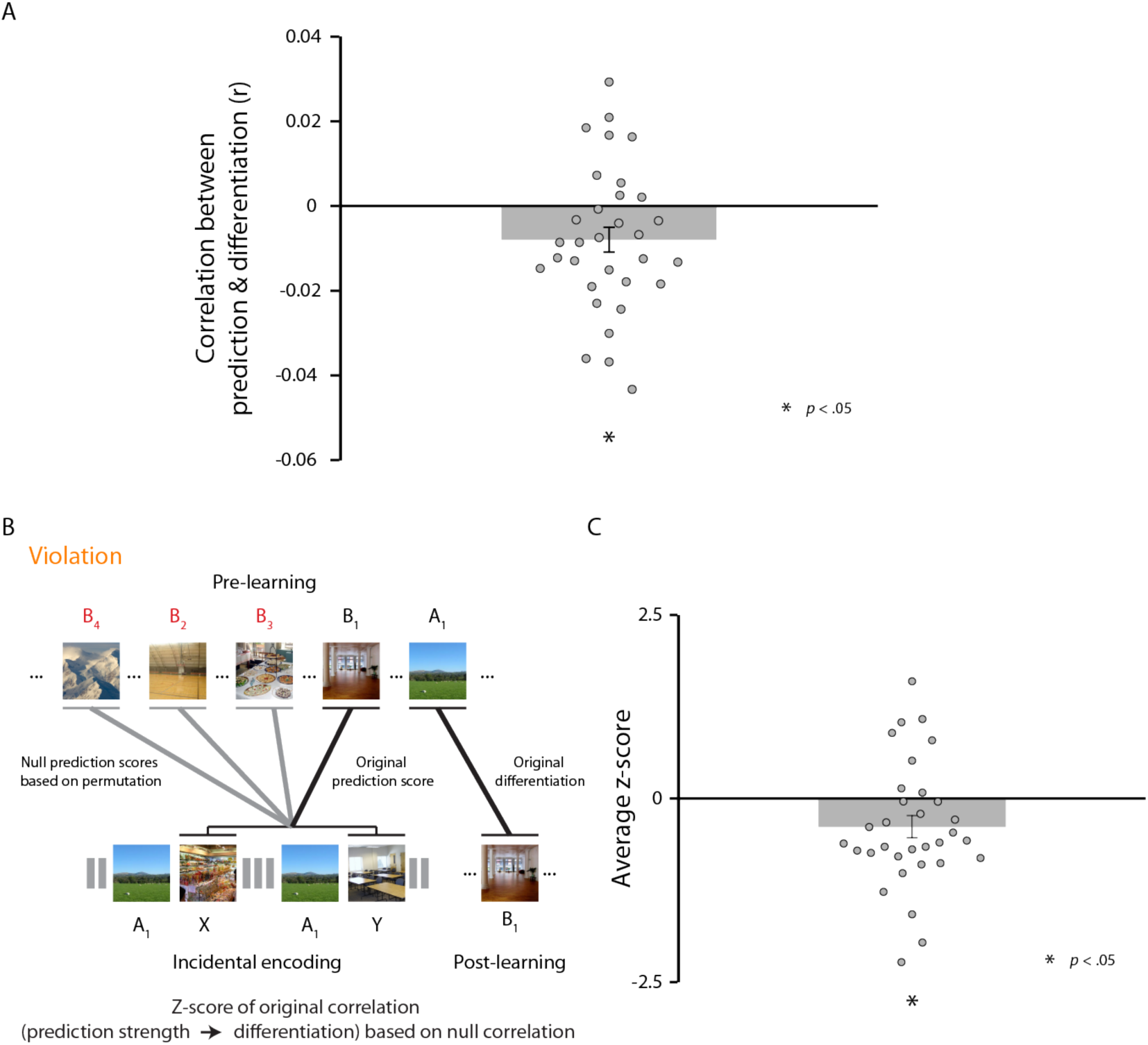
Relationship between prediction strength and differentiation, and item-specificity test for this relationship. (A) The relationship between prediction strength and differentiation was significantly negative. (B) Schematic of procedure for permutation analysis. For each participant, within the violation condition, we permuted the pairings of the pre-learning B snapshots and the violation (X/Y) trials for the associated A items 1,000 times. We then re-calculated the prediction scores and correlated them with the actual differentiation scores. Finally, we calculated a *z*-score for the original relationship with respect to the null distribution of relationships and assessed its reliability across participants. (C) The original relationship between prediction strength and differentiation was significantly more negative than the relationships based on permuted pairs.

### Ruling out univariate confounds

We have assumed that pattern similarity between A and B reflects a change in the relationship between the distributed representations of these items. However, univariate activation can affect pattern similarity (Coutanche, 2013; Davis and Poldrack, 2013; Davis et al., 2014; Aly and Turk-Browne, 2016), which could in principle explain some of our results. For example, weakened memory of B items in the violation condition might be expressed as lower activation in the post-learning phase, which could reduce pattern similarity for these items. This could also potentially explain the observed negative relationship between prediction strength and differentiation: greater misprediction could lead to more weakening, which (due to univariate confounds) could show up as lower pattern similarity. These scenarios are unlikely, however, in light of the item-specificity of the differentiation effect: If the differentiation effect were merely due to reduced activation of B items in the violation condition, then the same pattern of results should have persisted even after permuting the AB pairings.

Several additional results provide further evidence against this alternative account. First, univariate activation in left CA2/3/DG did not differ between the violation and non-violation conditions during the post-learning phase (*t*(31) = −0.55, *p* = 0.58). Second, there was no trial-by-trial relationship between univariate activation in the post-learning phase and differentiation (*t*(31) = 0.37, *p* = 0.71). Third, the negative relationship between prediction strength and differentiation persisted after controlling for the univariate activation level (during the post-learning snapshot phase) with partial correlation (*t*(31) = −2.63, *p* = 0.01). These observations are consistent with our interpretation that learning reflects differentiation of the underlying neural patterns rather than a change in overall activity.

## Discussion

The results of this study extend our prior work showing that mispredicted memories are weakened (Kim et al., 2014). Here we show that restudying a previously mispredicted item leads to differentiation of its hippocampal representation away from the prior context. We interpret this finding in terms of the nonmonotonic plasticity hypothesis (NMPH), which posits a U-shaped relationship between memory activation and learning: Low activation has no effect, moderate activation leads to memory weakening, and high activation leads to memory strengthening (Norman et al., 2006, 2007). Based on our prior study exploring effects of misprediction (Kim et al., 2014), we hypothesized that violation trials (where A was not followed by B) would elicit low-to-moderate levels of activation of the mispredicted B item, thereby weakening the synaptic connections between the (strongly activated) A item and the (moderately activated) B item. If memory is tested at this point, then the model predicts worse memory for the B item due to these weakened connections, as observed in Kim et al. (2014). Here, we explored what happens when the (weakened) B item is subsequently restudied. In this situation, our theory predicts that, due to the weakened connections to the shared features of A and B, activation spreads to new features that were not previously shared with A. These newly activated features at restudy are subsequently incorporated into the representation of B. This process of weakening connections to features (formerly) shared with A on the misprediction trial, and strengthening connections to features *not* formerly shared with A (on the restudy trial) has the net result of moving B’s representation away from A’s representation, thereby differentiating these patterns.

Our theory generates two additional hypotheses that we were able to test in the current study. First, neural differentiation should be competition-dependent, with stronger (but still moderate) prediction leading to more competition and greater subsequent differentiation. This hypothesis was supported by the observed negative relationship between prediction strength (at violation trials) and neural differentiation in left CA2/3/DG. Second, neural differentiation should occur with respect to the specific item that wins the competition and not other items. This hypothesis was supported by our randomization results, which showed that differentiation depends on the prediction of the specific B item that was previously paired with A, and that differentiation reflects the neural representation of B specifically moving away from the neural representation of A, as opposed to becoming uniformly more distinct from other scenes.

The NMPH implies that there will be boundary conditions on our conclusions. In particular, the pattern of results observed here — an overall differentiation effect, with greater prediction associated with more differentiation — will occur when activation for mispredicted items is in the low-to-moderate range. Because the NMPH posits that high activation leads to strengthening, much stronger predictions during violation trials may *strengthen* — rather than weaken — connections between A and unique features of B. This, in turn, will allow A to activate formerly unique features of B, leading to integration (i.e., increased neural overlap of A and B) instead of differentiation. We plan to test this prediction in future work by increasing associative strength between paired items (e.g., via more extensive exposure) or by adopting a more explicit prediction task (as opposed to the incidental approached used here).

A few other recent studies have used a similar pre-post “snapshot” approach to study differentiation (Schapiro et al., 2012; Hulbert & Norman, 2015; Favila & Kuhl, 2016; Schlichting et al., 2015). Schapiro et al. (2012) did so in a statistical learning paradigm, in which the transition probabilities between items varied: in the strong pair condition, A was always followed by B (transition probability = 1); in the weak pair condition, A was sometimes followed by B (0.33); and in the shuffled pair condition, A almost never was followed by B (~0). In CA2/3/DG, from pre- to post-learning phases (in which items were presented randomly), members of strong pairs showed increased neural overlap (integration) and members of weak pairs showed decreased overlap (differentiation), both relative to shuffled pairs. Hulbert and Norman (2015) used a retrieval practice paradigm with highly similar pictures of animals: Rp+ items were practiced, which should lead related Rp- items to activate as competitors; then Rp- items were restudied. The degree of left hippocampal differentiation predicted subsequent cued recall memory success. Favila and Kuhl (2016) showed that linking two scene stimuli to a shared face associate leads to hippocampal differentiation of the scenes. Lastly, Schlichting et al. (2015) used a similar paradigm exploring the effect of linking two unrelated objects to a shared associate. They looked at a wide range of regions and showed differentiation in some regions and integration in others. They also manipulated whether linked pairs were trained in a blocked (all AB prior to any BC) or interleaved (intermixed AB and BC) manner and found that integration was more prevalent after blocked training.

What is missing from these prior studies is a mechanistic explanation of why differentiation occurs, what determines the size of the effect, and when and where you get differentiation vs. integration. We hypothesize that the key mediating variable is the level of memory activation — moderate activation followed by restudy leads to differentiation, strong activation leads to integration. For example, in Schapiro et al. (2012), weak transition probabilities (0.33) during AB pair learning may induce moderate levels of prediction of B, leading to differentiation of A and B. In contrast, higher levels of prediction in the strong pair condition may lead to the opposite integration effect. Schlichting et al. (2015) also provides hints regarding our hypothesis. Specifically, blocked AB study may increase competitor activation during BC learning, tilting the balance towards integration. Likewise, regional differences in differentiation vs. integration may relate to how tightly activity is controlled: In regions like the hippocampus that have sparse activation, it is harder for related memories to activate strongly, biasing learning toward differentiation; other regions, including in cortex, with less sparse activation would be biased toward integration. Crucially, none of the above studies measured competitor activation, so they could not test our hypothesis. The main added value of our study is thus that we establish a link between competitive dynamics during learning and subsequent representational change.

Our finding of neural differentiation in left CA2/3/DG is distinct from the notion of hippocampal pattern separation (see Hulbert & Norman, 2015). Pattern separation refers to the fact that the hippocampus automatically assigns distinct representations to stimuli due to sparse coding in DG and CA3 (Yassa & Stark, 2011). While this pattern separation process reduces neural overlap in the hippocampus, there is still some residual overlap between similar items, which can lead to interference (Norman et al., 2003, 2005). Our differentiation mechanism operates on this residual overlap after standard pattern separation takes place. That is, the residual neural overlap between related memories of A and B leads to incorrect prediction of B when A is presented, which drives further reduction of the residual neural overlap and reduces subsequent interference.

This study was not designed to identify the behavioral consequences of neural differentiation, although that remains an important goal of future work. Here we used an item recognition memory test to be consistent with our prior study (Kim et al., 2014), but this is not a sensitive way to measure differentiation. Prior modeling work (e.g., Norman & O’Reilly, 2003; Norman, 2010) and empirical studies (e.g., LaRocque et al., 2013) suggest that reduced neural overlap can have opposite effects on different components of memory: boosting *recollection* by reducing interference from other memories, but reducing *familiarity* by lowering global match. Because these effects go in opposite directions, they might cancel each other out in our recognition test, which is sensitive to both components. The aforementioned study by Favila and Kuhl (2016) suggests a better way of behaviorally measuring differentiation: They found that neural differentiation between visually-similar scenes (e.g., A and B) led to enhanced subsequent learning of new associations between these scenes and objects (A-X and B-Y), presumably because of reduced interference between the scenes.

### Summary

We found that interleaved misprediction and restudy leads to neural differentiation. These findings are in line with predictions from our neural network modeling work (Norman et al., 2006, 2007) and other recent studies in the field (Schapiro et al., 2012; Favila and Kuhl, 2016; Hulbert and Norman, 2015; Schlichting et al., 2015). Most importantly, by revealing a relationship between prediction strength and differentiation, this work suggests a key role for competition in driving representational change. This complements prior results showing that activation of non-target memories can weaken them (Detre et al., 2013; Kim et al., 2014; Newman & Norman, 2010; Poppenk and Norman, 2014), and suggests that one function of such weakening is to prepare the memory to accept new features and associations. This adaptive optimization of memory may serve to increase the accuracy of memory with respect to the current environment, to reduce subsequent interference, and to minimize prediction errors.

## Acknowledgements

NIH R01 MH069456 to KAN and NBTB, R01 EY021755 to NBTB

